# RNA-seq derived sequence variations are excellent features for cell line identification

**DOI:** 10.1101/2025.08.22.671765

**Authors:** Lisa Müller, Simon Müller, Khursheed Ul Islam Mir, Jana Lange, Sven Hagemann, Alice Wedler, Frank Hause, Claudia Misiak, Danny Misiak, Tony Gutschner, Stefan Hüttelmaier, Markus Glaß

## Abstract

Cell lines are indispensable models for analyzing molecular mechanisms underlying human diseases. However, incorrect annotation and cross-contamination can introduce severe bias in respective studies. Accordingly, various publishers request authentication of cell lines before publication. Short tandem repeat profiling is commonly used to verify cell line identity and purity but does not guarantee that published results are based on the samples tested by this method. In this study, we demonstrate that RNA-seq-derived sequence variation information is eligible for unambiguous cell line-specific clustering. Based on this finding, we propose methods for reliable cell line identification from RNA-seq data using supervised machine learning methods. In addition, we demonstrate the ability to detect cross-contamination of human cell lines. The presented methods are insensitive to different data pre-processing steps and quality measures. The proposed TopFracCCLE algorithm for cell line identification and detection of cross-contamination is available as R-script at https://github.com/HuettelmaierLab/topFracCCLE.

## 2 Introduction

Cell lines have been used for decades as models for human diseases as well as for virus production, vaccine development and for synthesizing recombinant proteins [1]. However, mis-labeling and cross-contamination are frequently observed in research labs, requiring the occasional re-authentication of cell models, especially before publishing. Current estimates suggest that up to 36 % of cell lines are cross-contaminated or misidentified [2], indicating the necessity for rapid, cost-effective and easy-accessible methods for cell line authentication.

Cell line models are derived from individual patients or animals and thus carry the unique genomic code of the respective donor. Service providers offer authentication, usually based on short tandem repeat (STR) profiling. This method has been recommended by the American Type Culture Collection (ATCC) [3] and is frequently requested to prove the identity of cell lines used in a specific experiment. However, STR profiling does not guarantee that published results have been generated with the cells being submitted for testing, since interchanges or contamination may occur after performing the respective tests. Next generation sequencing (NGS) data provide excellent options for cell line authentication, since they offer DNA/RNA sequence information at high resolution, allowing the rapid identification of sequence variations. Although high-throughput RNA sequencing (RNA-seq) provides less variant information than DNA-based whole genome sequencing (WGS), it has been shown that RNA-seq-derived sequence variations can be used for cell line identification [4, 5].

Here, we consolidate and extend this finding by exploring requirements and limitations for robust cell line identifications based on RNA-seq-derived sequence variation information. We used unsupervised clustering to demonstrate that RNA-seq samples from distinct cell lines form separate clusters, even at comparatively moderate sequence coverage. We have trained a k-Nearest-Neighbor (kNN) classifier with sequence variation data from 22 distinct cell line models. Classification yielded perfect classification results for our in-house as well as publicly available datasets. Exploiting RNA-seq information of more than 1700 cell lines, provided by the cancer cell line encyclopedia (CCLE, [6]) project, we finally implemented a method for human cell line identification that, in addition, can be used for detecting cross-contamination of human cell lines.

## 3 Materials and methods

### 3.1 Cell culture

Cells were cultured in the appropriate medium supplemented with 1 % GlutaMAX (Thermo Fisher Scientific, Waltham, MA, USA) and fetal bovine serum (FBS) at the appropiate concentration at 37 C and 5 % CO2. Detailed information are provided in Table S1.

### 3.2 Library preparation and RNA sequencing

RNA integrity was assessed using a Bioanalyzer 2100 (Agilent, Santa Clara, CA, USA). RNA sequencing (RNA-seq) libraries were prepared according to the manufacturer’s instructions. For mRNA-seq, total RNA was used as input for poly (A) enrichment with oligo (dT) beads. After fragmentation, the first strand cDNA was synthesized using random hexamer primers. Then, the second strand cDNA was synthesized using dUTP, instead of dTTP. The directional library was ready after end repair, A-tailing, adapter ligation, size selection, USER enzyme digestion, amplification, and purification. For small RNA-seq, 3’ and 5’ adaptors were ligated to 3’ and 5’ end of small RNA, respectively. Then, the first strand cDNA was synthesized after hybridization with reverse transcription primer. The double-stranded cDNA library was generated through PCR enrichment. After purification and size selection, libraries with insertions between 18 and 40 bp were ready for sequencing. Library preparation and sequencing were performed either by Novogene (Munich, Germany) or in-house (Details in Table S2.)

Sample preparation of SW1990 and Panc-03.27 for cell line authentication and RNA sequencing was performed in parallel. Cells were trypsinized and counted to ensure 1.5x10^6^ cells in each sample. For both cell lines, non-mix-up samples contained 100 % of SW1990 or Panc-03.27 cells, respectively. For mix-up samples, three conditions were performed: 50 % of SW 1990 + 50 % of Panc-03.27 cells, 25 % of SW1990 + 75 % of Panc-03.27 cells, and 10 % of SW1990 + 90 % of Panc 03.27 cells. All samples were run in triplicates. For RNA sequencing, cells were homogenized in Trizol and total RNA isolated by phenol/chloroform extraction and isopropanol precipitation. Library preparation (Poly(A) tail RNA selection) and RNA sequencing (total RNA amount ≥ 0.2 *μ*g) was performed by Novogene (Munich, Germany) on a NovaSeq X Plus Series system (Illumina, San Diego, USA).

### 3.3 STR profiling

Cell lines were authenticated by Eurofins Genomics (Ebersberg, Germany) STR profiling service, which carried out DNA isolation from cell pellet and determined genetic characteristics by PCR-single-locus-technology using the AmpFlSTRIdentifilerPlus PCR Amplification Kit (Thermo Fisher). This kit comprised 15 core loci: D8S1179, D21S11, D7S820, CSF1PO, D3S1358, tyrosine hydroxylase (TH01), D13S317, D16S539, D2S1338, D19S433, von Willebrand factor (vWA), human thyroid peroxidase gene (TPOX), D18S51, D5S818, and *α*-Fibrinogen (FGA) plus the gender-determining marker amelogenin (AMEL). Based on electropherograms, the analytical report lists all 16 loci and calls (cf. tables S9, S10). These alphanumerically scored alleles were entered into the STR profile database for human cell lines of the German Collection of Microorganisms and Cell Cultures (DSMZ) ([7, 8]) to compare to reference databases by checking for percent match between cell lines or mix-ups.

### 3.4 Pre-processing of RNA-seq data

Adapter sequences as well as low quality read ends were clipped off using Cutadapt. The processed sequencing reads were aligned to the human reference genome (UCSC hg38) using HiSat2 [9]. Samtools [10] was used to extract primary alignments and to index the resulting BAM files. FeatureCounts [11] was used for summarizing gene-mapped reads, using ENSEMBL v100 [12] as annotation basis. Down-sampling of BAM files was performed using the sample command from the bedtools suite [13].

### 3.5 Generation of simulated cell line contamination data

To simulate contamination of cells from two distinct human cell lines, defined numbers of sequencing reads were randomly sampled from fastq files of pure samples using the seqtk sample function (https://github.com/lh3/seqtk). Subsequently, reads from the different cell lines were merged together in new fastq files and processed as described above.

### 3.6 Variant calling

Variant calling from alignment files in BAM format was performed using mpileup and call commands from the bcftools suite [14]. The human UCSC hg38 genome sequence was used as reference. Base calling quality values of at least 20 and sequencing depths of at least 10 reads were used as filter criteria.

### 3.7 Clustering and k-nearest-neighbor classification

Variants found in each sample were collected in a binary matrix assigning the presence of all found variants in each sample. Variants found exclusively in one sample were removed from the matrix and pair-wise Jaccard-distances of all samples were calculated using the remaining variants. Clustering was performed using the hclust function of R [15] based on the distance calculations and application of the ward.D2 algorithm. Down-sampling was performed by random sampling of aligned reads from bam files using the sample function of the bedtools suite [13].

For kNN training, the knn3Train function of the caret R-package [16] was applied using the variant-presence matrix.

### 3.8 CCLE data acquisition and TopFrac classification

CCLE cell line single nucleotide polymorphism (SNP) data were gathered from depmap via the ExperimentHub R-package, table mutationCalls 22Q2 [17]. These data were used to construct a binary matrix specifying the presence of all found SNPs in each cell line. Since the depmap-provided data annotations were based on the human genome build GRCh37, genomic positions were updated to GRCh38 using the UCSC liftover website (https://genome.ucsc.edu/cgi-bin/hgLiftOver).

For cell line identification, the number of common SNPs from the query sample and each cell line was determined.

### 3.9 Figures

All figures were created using the R-packages ggplot2 [18] and patchwork, if not explicitly stated otherwise.

## 4 Results

### 4.1 RNA-seq-derived variant calling leads to cell line-specific clustering

For assessing if RNA-seq samples from multiple distinct cell lines form homogeneous clusters, we assembled a dataset comprising 90 poly-A enriched samples from 22 distinct cell lines containing at least three samples per line. These samples represent control experiments from various experiments performed in our lab and comprise either wild type cells or cells transfected with control siRNAs and have been sequenced between 2017 and 2024. Cell culture, RNA isolation and library preparation have been performed by distinct experimenters and sequencing has been conducted by different facilities (cf. Table S2). Thus, the samples displayed a high degree of heterogeneity in respect to processing and technological settings. Variant calling, using the human genome (UCSC hg38) as a reference, revealed a high variability in the number of detected variants, as expected. These numbers not only drastically differ between distinct cell lines but also show substantial inter-experimental batch variation, in part even between experimental replicates (Fig. S1A).

One important factor influencing the number of variants detected is sequencing depth. This is reflected by a strong positive correlation between the number of sequenced reads and the number of variants detected for the respective sample (Fig. 1A, S1). For all samples, the vast majority of detected sequence variations was unique to the respective sample (Fig. 1B). Since cell line clustering would not benefit from including variations only present in one particular sample, sample-specific variants were ignored while constructing a binary matrix assigning all detected variants shared by at least two samples to these respective samples. Furthermore, the removal of sample-exclusive variants reduced storage amount and runtime to create the assignment matrix considerably. Hierarchical clustering using Jaccard distances between the samples yielded sub-trees consisting entirely of samples from the same cell lines. After sub-dividing the obtained dendrogram into the 22 top sub-clusters, each sub-cluster contained samples from exactly one cell line and each cell line was represented by exactly one sub-cluster (Fig. 1C, Table S3). Thus, RNA-seq derived variations from a reference genome sequence are suitable for the discrimination of samples belonging to a certain cell line model. Notably, experiment-specific samples from the same parental cell lines formed sub-clusters as well, indicating that even passage- or experimentally motivated sequence variations are identified (Fig. 1C, Table S3).

**Figure 1:**
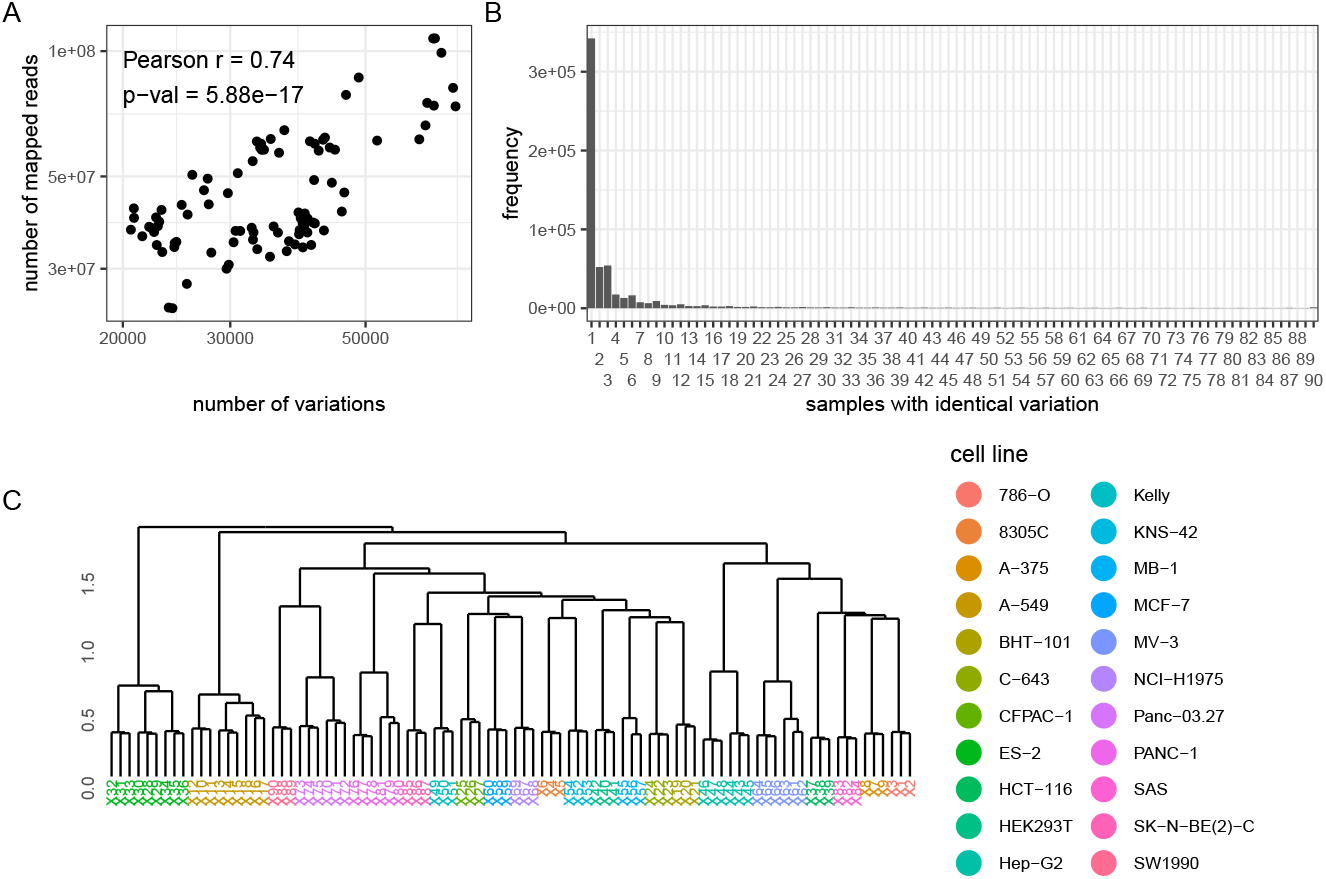
Clustering of cell lines by poly-A enriched RNA-seq-derived sequence variations. **A** Relation between detected sequence variations and mapped reads. **B** Histogram showing the number of variations shared by a certain amount of samples. **C** Hierarchical clustering of samples based on sequence variations.

For comparison, we also performed hierarchical clustering using normalized expression values (FPKM) of all protein-coding genes as input. However, although samples derived from the same experiments clustered together, accurate clustering of cells of the same origin was not achieved with this approach. For example, samples from the pancreatic cancer-derived cell line PANC-1 (samples X76 - X81) were distributed in very distinct sub-clusters. Subdividing the dendrogram into the 22 top clusters revealed further discrepancies in the cluster identities (Fig. S2). This indicated that, without further feature-selection procedures, the binary representation of sequence variants is a more robust measure for clustering samples according to cell line identity.

Since the number of detected variants depends on sequencing depth, we investigated how many sequencing reads are sufficient to retain optimal clustering results. Thus, we randomly sub-sampled the reads to obtain pre-defined numbers, performed hierarchical clustering as described above and divided the resulting dendrograms into 22 clusters and examined (a) if these clusters contain exclusively samples from one cell line and (b) if all samples belonging to the same cell line were included in exactly one cluster. As expected, the number of detected variants decreased with reduced sequencing depth (Fig. S3A). However, these investigations revealed that as few as 500,000 reads per sample were sufficient to optimally cluster the samples according to the cell line model (Fig. S3B). Thus, since typical RNA-seq samples, e.g. created for assessing differential gene expression, usually comprise more than 10 million reads or read-pairs, sequencing depth seems not to be a limitation for reliable cell type-specific clustering in a common setup.

To next test, whether the described sequence variation-based clustering approach is capable of discriminating cell line identities from microRNA (miRNA) expression profiles as well, we assembled another data set consisting of 18 small RNA-seq samples representing six distinct cell lines. Sequencing read numbers mapped to the human reference genome ranged from 11.4 million reads to 30.1 million reads per sample.

Sequence variations found in these samples were significantly fewer in numbers compared to the poly-A enriched samples, ranging between 151 (MB-1) and 5194 (A-549). Yet, these low numbers were sufficient to reliably cluster the samples according to their cell line identities (Fig. 2A,B).

**Figure 2:**
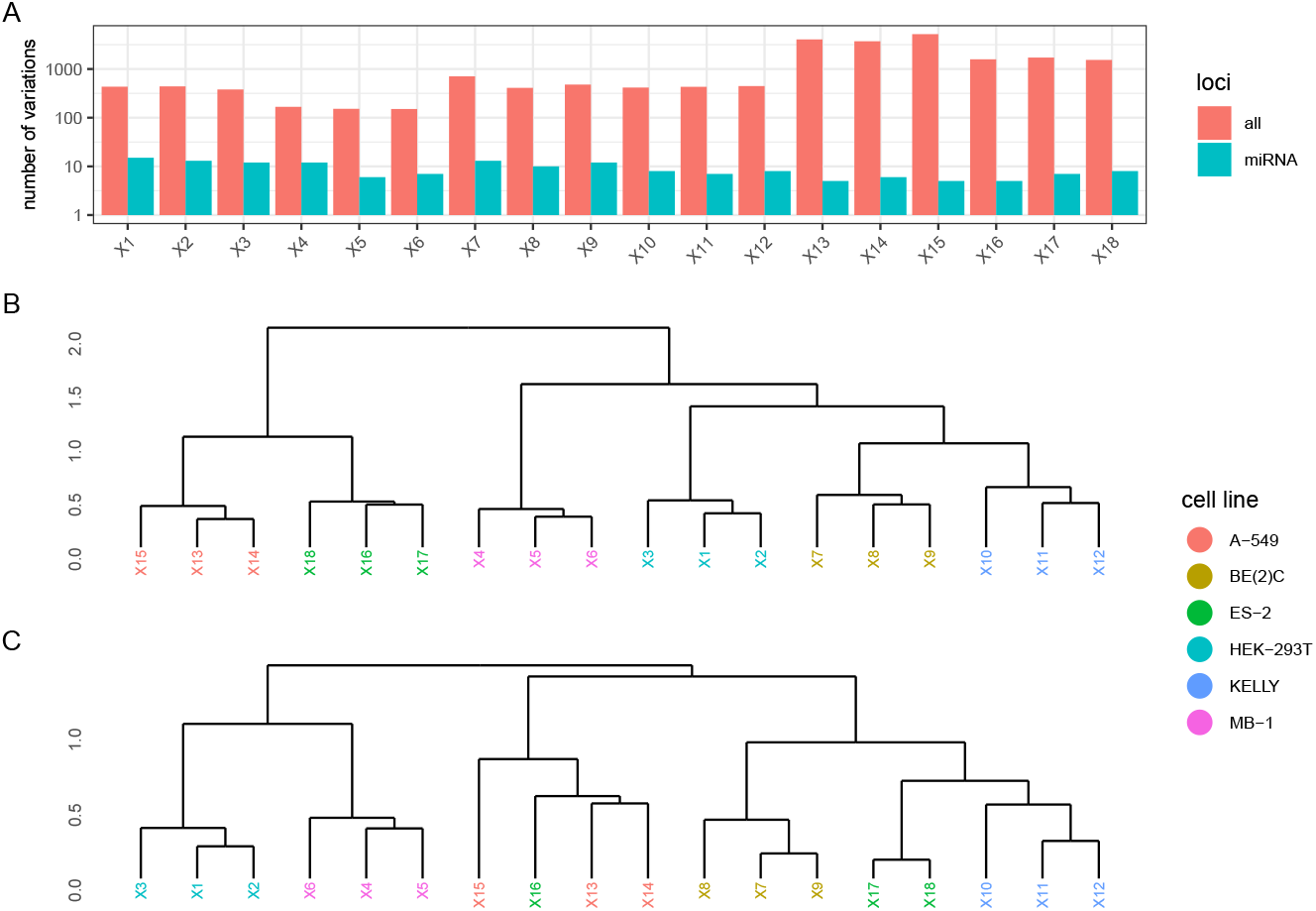
Clustering of cell lines by small RNA-seq-derived sequence variations. **A** Numbers of sequence variations found in the whole sample and miRNA loci. **B** Hierarchical clustering using all detected, non sample-exclusive, variations. **C** Hierarchical clustering using only non sample-exclusive variations detected in miRNA loci.

Small RNA-seq libraries usually are generated by selecting short PCR products and, thus, contain cDNAs of various short RNA types, including miRNAs. Accordingly, the majority of variants detected in small RNA-seq samples were found in loci not covering miRNA transcripts. However, restricting detected variants to those located in known miRNA loci, according to miRBase v22 [19], reduced the number of variants per sample to less than 20, leading to sub-optimal clustering results (Fig. 2C). Thus, although our results demonstrate the feasibility to differentiate cell lines by utilizing sequence variations obtained from small RNA-seq samples, variation amounts found inside miRNAs loci alone were not sufficient to unambiguously separate the samples by cell line identity.

### 4.2 Nearest-Neighbor classification leads to reliable prediction results

Since hierarchical clustering based on sequence variations can be applied to distinguish samples from different cell line models reliably, we hypothesized that sequence variation information can be used to accurately predict cell line identities from unknown sources as well. To test this hypothesis, we implemented a kNN trained on a new dataset (in-house_polyA_) containing exactly three samples per cell line. Thus, the set contained 66 samples from 22 cell lines. Analogous to the hierarchical clustering, the presence of distinct sequence variations was used as feature vector. Sample-exclusive variants were not considered, in order to reduce the feature-space.

For assessing the performance of the classifier, we conducted a balanced three-fold cross validation. I.e., in each of three iterations, we used two thirds of the samples from our data set for training and the remaining third for testing the kNN (k = 2). The training sets contained two, the test sets one sample per cell line. In each iteration of the cross validation, all cell line identities of the test set were predicted correctly, yielding an overall prediction accuracy of 100 %. To test generalization capability of our kNN on completely unrelated samples, we assembled validation sets from publicly available alignment files (.bam) obtained from the ENCODE project [20]. The first validation set, ENCODE_polyA_, contained 14 samples, generated from poly A-enriched libraries. It was comprised of 10 samples, two each from five distinct wild type cell lines (A-549, PANC-1, HCT-116, MCF-7, Hep-G2) as well as another four Hep-G2 samples, two transfected with shRNAs against SRSF1 and two transfected with shRNAs against DDX55 (cf. Table S4). The other validation set, ENCODE_total_, contained five samples from five distinct cell lines (A-375, Hep-G2, HCT-116, MCF-7, PANC-1, cf. Table S5), generated from total RNA libraries. Variant calling results of the samples from the validation sets were used as input for a kNN trained on the whole in-house_polyA_ dataset (66 samples), setting the number of the nearest neighbors parameter k to three. All samples of the two validation sets were predicted as annotated (Fig. 3).

**Figure 3:**
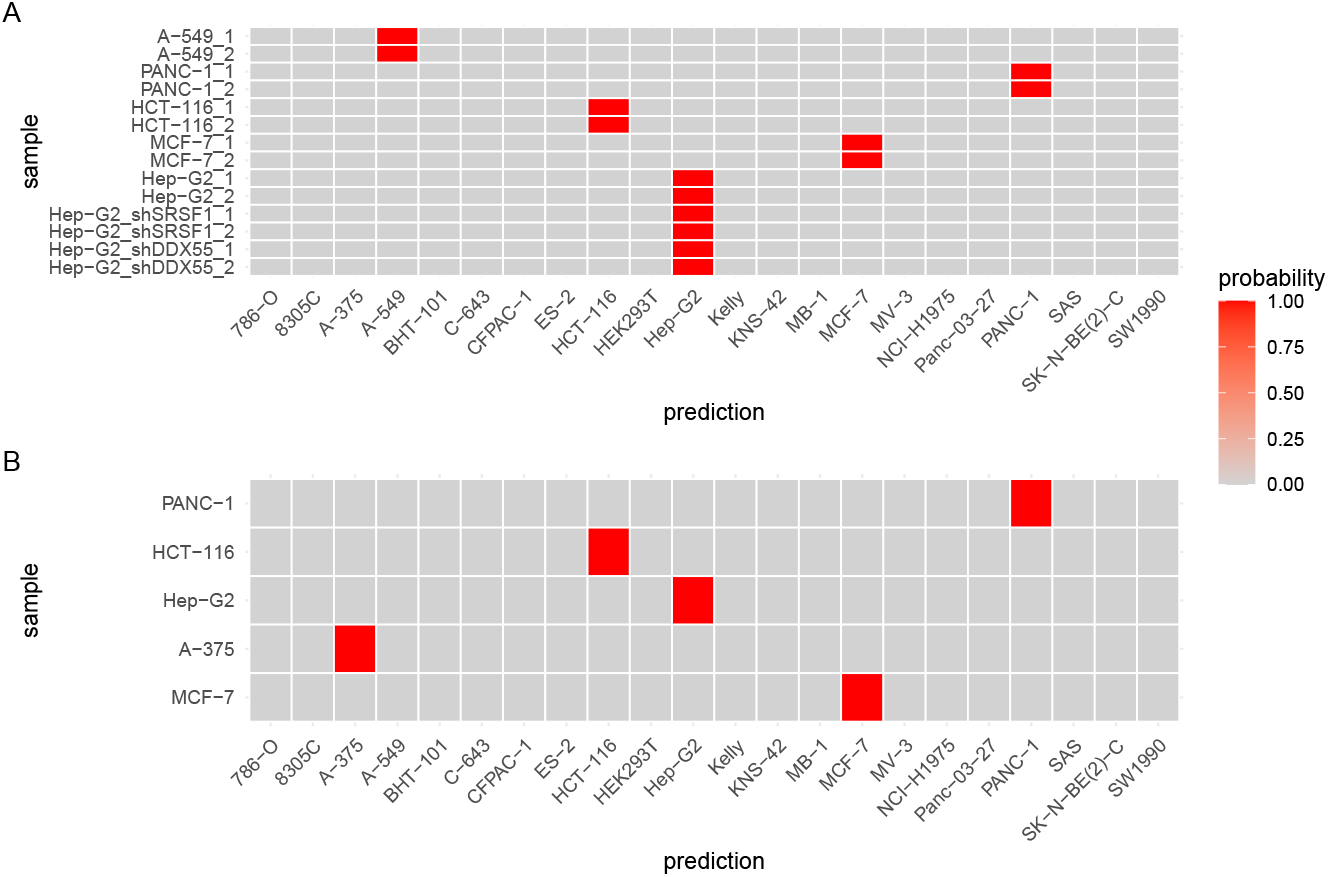
kNN classification results. **A** Prediction probabilities of samples from set ENCODE_polyA_, containing poly A-enriched RNA-seq samples. **B** Prediction probabilities of samples from set ENCODE_total_, containing total RNA-seq samples.

### 4.3 Variant comparisons are suitable to universally identify human cell lines and to detect cell line cross-contaminations

In the previous section, we demonstrated the feasibility to reliably determine the identities of cells from a small portfolio of in-house sequenced cell lines. However, in order to develop a method for the general determination of cell line identity from an unknown or uncertain sample, a more comprehensive resource was desirable. For this purpose we took advantage of sequence variant information provided by the cancer cell line encyclopedia (CCLE) project [6]. The CCLE data set, obtained via the ExperimentHub [17] resource, comprised SNP data of 1771 distinct human cell lines. In contrast to our previously described data sets, the CCLE data set was lacking replicate samples, i.e., there was only one feature vector per cell line available. Furthermore, the number of reported SNPs per individual cell line differed tremendously from 16 to 30821 (Fig. S4). These properties of the CCLE data set rendered it unsuitable for the previously proposed nearest-neighbor classification approach. Thus, to leverage the CCLE data set for cell line identification, we implemented a rather simple method comparing the SNPs found in the query sample with those provided by the CCLE data set. Prediction results are derived by sorting the comparison results by the relative abundance of SNPs found in a certain cell line. I.e, the cell line of which the highest percentage of SNPs matches is presented as the top candidate for cell line identity. Figure 4A shows a scheme of the method we called TopFracCCLE. To assess prediction accuracy, we predicted cell line identities for our in-house samples as well as the aforementioned publicly available data sets ENCODE_polyA_ and ENCODE_total_. However, since four of the cell lines from the set in-house_polyA_ were not contained in the CCLE data set (HEK-293-T, C-643, MV3 and SK-N-BE(2)-C), we excluded those lines and created the set in-house2_polyA_ comprised of triplicate samples from 18 distinct cell lines, totaling in 54 samples.

**Figure 4:**
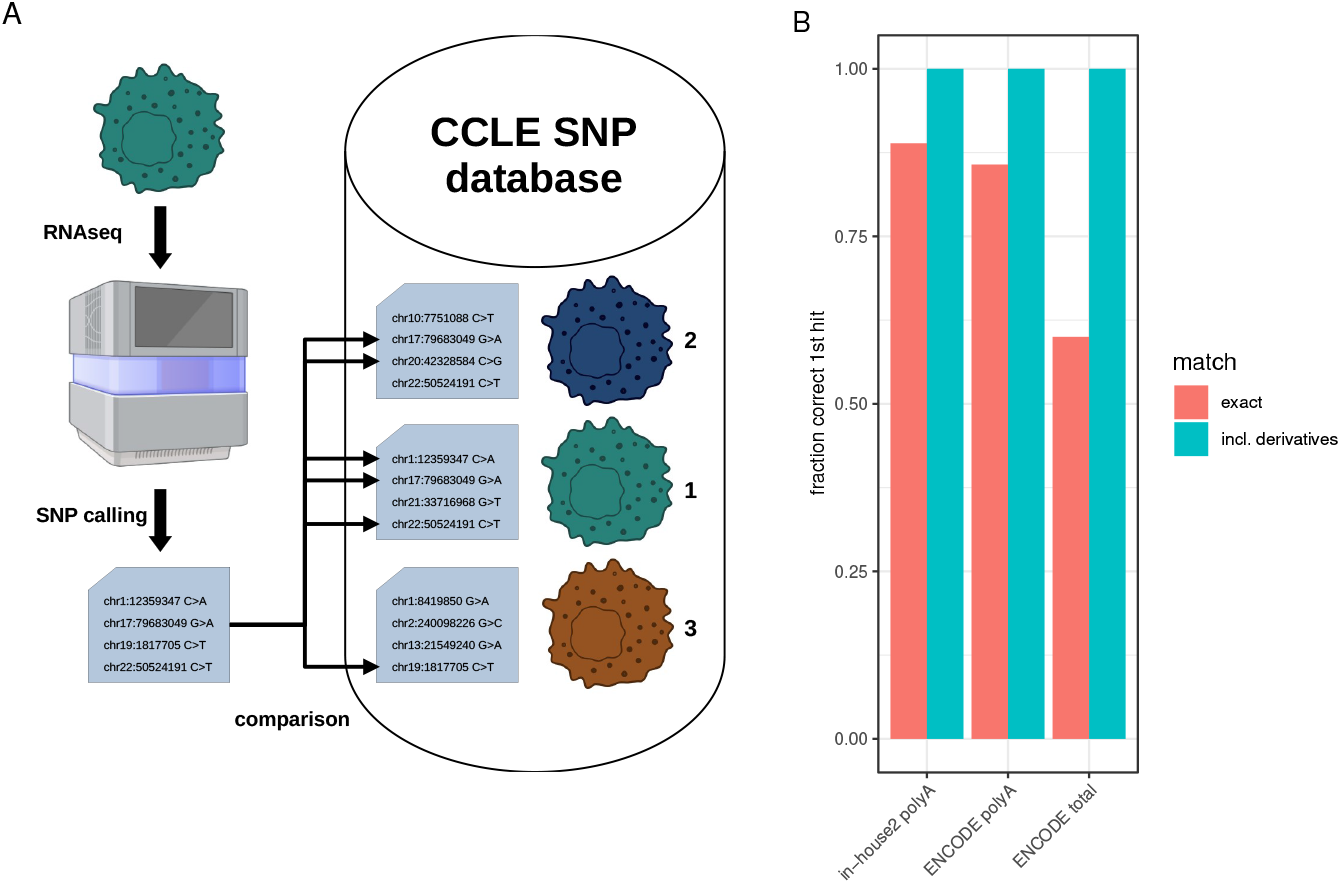
TopFracCCLE classification results. **A** Scheme for predicting cell line identities using CCLE-derived SNP data. Image in part created with BioRender. **B** Prediction results on three distinct test sets. Numbers refer to the fraction of correctly predicted cell lines reported as first hit, exactly matching the known query line or either the matching line or a line known to be contaminated with or a derivative of the query line.

For the majority of samples from all three validation sets, the cell line showing the highest fraction of SNP overlap (1st hit) was the expected cell line. Seemingly incorrect predictions were related to only two cell lines, A-375 and MCF-7. However, in all these cases, the reported first hit was a derivative of or was reported to be contaminated with the query cell line, according to CCLE recordings. I.e., the three cell lines matching best the samples of the melanoma-derived cell line A-375, regardless from which validation set, were A375 SKIN CJ1, A375 SKIN CJ2 and A375 SKIN CJ3, followed by the expected cell line (A-375) as the fourth best hit. These cell lines were derived from A-375 cells by selection for drug resistances [21, 22]. All samples of the breast cancer derived cell line MCF-7 were predicted as cell line KPL-1. According to Expasy Cellosaurus [22], this cell line is actually a MCF-7 derivative. Thus, if the contaminated/derived cell lines are considered correct as well, TopFracCCLE correctly predicted all samples from the three sets (Fig. 4B). Fractions of SNPs found in correct/derived cell lines ranged from nine to 48 percent and were considerably lower in other, non-derived cell lines (Tables S6-S8).

Since the TopFracCCLE approach is based on comparing SNPs found in a query sample with those reported for distinct cell lines, we hypothesized its applicability on detecting cross-contaminations of human cell lines. More precisely, if cells from two or more cell lines were present in a given sample, RNA sequencing data are expected to contain SNP information matching a substantial fraction of SNPs reported for the respective parental cell lines.

We tested this hypothesis by simulating contamination *in silico* and *in vitro*. For *in silico* contamination experiments, sequencing reads from wild type samples of lung adenocarcinoma-derived H-1975 cells and neuroblastoma-derived KELLY cells were sub-sampled and mixed at defined ratios. For *in vitro* simulation of cell line contamination, we mixed cells from the pancreatic adenocarcinoma (PDAC)-derived cell lines SW1990 and Panc-03.27 at defined ratios. Triplicate samples from each mixture ratio were sequenced and variants were detected. For comparison, DNA from the same samples was sent for STR profiling by Eurofins Genomics. Additionally, sequencing reads from pure SW1990 and Panc-03.27 cells were mixed *in silico* for comparisons. Regardless, if cells were mixed genuinely or *in silico*, the numbers of detected SNPs belonging to a certain cell line increased with the real/expected fraction of these cells in the respective samples (Fig. 5A,C, Fig. S5A). Whereas in pure KELLY, SW1990 and Panc-03.27 samples approximately 40 % of reported SNPs were detected, this proportion was approximately 30 % in H-1975 cells.

**Figure 5:**
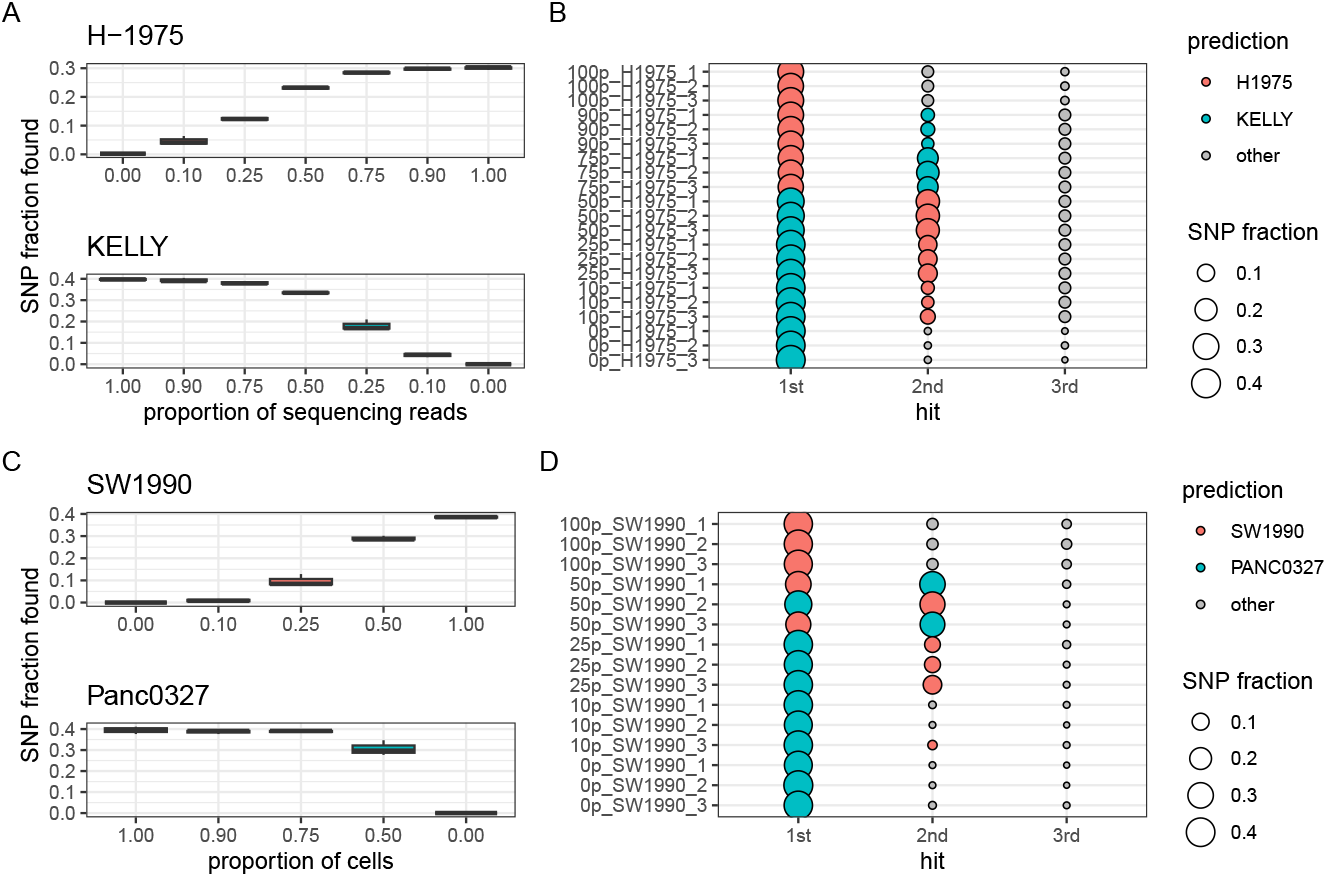
TopFracCCLE predictions on *in silico* and *in vitro* contamination simulations. **A** Proportion of detected SNPs associated with H-1975 or KELLY cells in dependence of the proportion of sequencing reads originating from samples of the respective cell line. **B** First, second and third hit cell line reported by TopFracCCLE for the respective mixture samples. Numbers in the sample labels refer to the percentage of reads from pure H-1975 samples. **C** Proportion of detected SNPs associated with SW1990 or Panc-03.27 cells in dependence of the proportion of cells from the respective line in the samples. **D** First, second and third hit cell line reported by TopFracCCLE for the respective mixture samples. Numbers in the sample labels refer to the percentage of SW1990 cells contained in the samples.

Samples containing a majority of cells/reads from one line got assigned this line as first hit by TopFracCCLE. For the H-1975/KELLY samples, 1:1 mixture of reads got consistently assigned KELLY as first hit and H-1975 as second hit. This suggests, that a higher SNP fraction detected in non-contaminated cell populations shifts prediction results towards the line with more SNPs detected when mixed at equal ratios. Accordingly, *in vitro* and *in silico* 1:1 mixtures of SW1990 and Panc-03.27 cells resulted in either the first or the second line to be predicted as first hit. In the *in silico* mixture of H-1975/KELLY cells, the lower concentrated cell line was obtained as second hit down to a fraction of 10 % of this line. In the SW1990/Panc-03.27 mixtures, the lower concentrated cell line was reliably found as second hit down to a fraction of 25 %, in both, *in silico* and *in vitro* simulations (Fig. 5B,D, Fig. S5B).

To compare the results of the TopFracCCLE method with the gold standard, we sent the *in vitro* mixed samples for STR profiling to Eurofins Genomics. Using non-empty markers as scoring mode and a minimum of 14 shared markers, the non-mixed-up samples showed 100 % similarity with SW1990 (samples CL00020652-654) and Panc-03.27 (samples CL00020655-657) cell line, respectively. The second best hits of these samples had similarities below 60 % (Table S10). In contrast, each mix-up sample (CL00020658-666) showed highest similarity with Panc-03.27 (89.3 %) and the second highest with SW1990 (75 %) cells (Table S10). These numbers align well with conclusions of the ANSI-ATCC ASN-0002 standard, suggesting that a similarity score below 70 % is indicative for unrelated donors and similarity scores between 70 % and 79 % are obtained either due to significant genetic drift or a mixture of two or more cell lines [1]. Further, since more than two alleles at a locus are rarely seen in normal human DNA profiles [1], multiple peaks at one locus indicate mixed populations of cells. With respect to a consensus profile, the six non-mixed samples (CL00020652-657) contained two alleles at each locus, which is in line with 100 % similarity of these samples to the SW1990 or Panc-03.27 cell line, respectively. More importantly, all nine mix-up samples (CL00020658-666) contained an additional allele at five of 16 loci as well as two additional alleles at one of these loci (Table S9) indicating cross-contamination in these samples.

In summary, these results suggest that the proposed TopFracCCLE method is suitable for detecting cross-contamination of two human cell lines, provided that at least 25 % of cells belong to the lower concentrated cell line. Based on our findings, we suggest to consider detected SNP fractions above 5% as a strong indicator for putative presence of a certain cell line model in a given sample.

## 5 Discussion

Mis-classified and cross-contaminated cell lines lead to incorrect conclusions, prevent reproducibility and unnecessarily waste resources. Cell line authentication is therefore crucial to ensure integrity of scientific results.

In the present study, we demonstrate the applicability of RNA-seq-derived sequence variations for robust cell line identification and detection of cross-contamination. We demonstrate that basic machine learning algorithms are sufficient to reliably discriminate between a multitude of distinct cell line models.

Hierarchical clustering based on the presence of detected sequence variations yielded cell type specific clusters. Furthermore, samples containing the same cell line models but sequenced for different studies were also distinguishable with this approach. This indicates that lineage divergence, occurring through passaging, could be monitored by this approach. Furthermore, although not directly related to cell line identification, it suggests the possibility of other applications of RNA-seq derived sequence variations, e.g., longitudinal tracking of genomic changes caused by reagents or factors influencing genomic stability or DNA damage repair mechanisms. Clustering results revealed to be insensitive to different pre-processing steps and varying sequencing depths. Notably, we further demonstrated the possibility to discriminate cell lines utilizing sequence variations derived from small RNA-seq data, despite far less detected variants in these samples.

Using a kNN approach we were able to predict cell line identities from a panel of 22 distinct cell lines, using RNA-seq samples either gathered in-house or from public sources as input. The method can be easily adapted to other cell line panels derived from arbitrary organisms. However, the applied kNN algorithm is rather resource-intense when applied to larger training data. Machine learning algorithms that generate class representatives instead of using the complete training data for prediction, e.g., k-means or support vector machines, should be appropriate alternatives reducing run time and storage demands.

Our TopFracCCLE approach reports for a given set of sequence variations the cell line that best matches these variants from a repertoire of more than 1700 human cell line models. We observed that for few of the tested cell lines the best data base hit was a cell line model derived or contaminated with the actual cell line. However, in these cases the fraction of detected sequence variations associated with the correct cell line was always close to the numbers determined for the derivatives/contaminated cell lines. Actually, this finding underlines the need for proper cell line identification procedures, since former mis-classifications led to incorrect information content, e.g., in public databases.

Furthermore, we successfully applied this algorithm to detect simulated cross-contaminations of human cell lines *in silico* and *in vitro*. Contamination of at least 25 % with another human cell line resulted in both cell lines appearing as the two best hits for the tested cell lines. This seems to be less sensitive than STR profiling, since with this DNA-based approach, all tested contamination ratios yielded the same similarity scores above 70 % for both contained cell lines, proving contamination also at lower levels. However, our method provides distinct advantages compared to STR profiling. Since our method is applied after sequencing, results refer to the actually processed samples. Furthermore, testing cell line identities using TopFracCCLE is not associated with further costs, once RNA-seq has been performed. And finally, the proportion of detected SNPs is correlated with the proportion of a particular cell line’s presence. Consequently, although the actual fraction of variants detected by TopFracCCLE varies with each cell line and the relationship between cell line abundance and detected SNP proportion seems to be non-linear, estimations about the actual proportions of the present cell lines seem possible. Such quantitative estimations regarding contamination extend are not possible to be derived from STR profiling.

Our described variant calling approach yields several thousand SNPs per sample. The median number of SNPs per cell line reported in the CCLE data set was 338. Although we did not observe this problem with the cell lines we tested, we expect that cell lines associated with a very limited set of reported variations might be hard to classify correctly. For example, cell line OUMS-27 is associated with only 16 SNPs in the CCLE data set. However, PANC-1 cells, associated with 35 SNPs, have been correctly identified in all of our test sets.

In summary, our study highlights the potential of RNA-seq derived sequence variations for cell line identification and detection of cross-contamination. We present practical implementations for these tasks yielding reliable results for a multitude of different cell line models.

## 7 Acknowledgments

We thank the ENCODE project and the labs of Thomas Gingeras, Richard Myers, Barbara Wold and Brenton Graveley for providing publicly available RNA-seq data via the ENCODE portal.

## Author contributions

*LM and MG conceptualized the study and wrote the paper. MG conducted data analyses. LM, SM, KUIM, JL, SHa, AW, FH and CM conducted the experiments. DM supported NGS data analyses. SHü and TG provided overall supervision and support. All authors reviewed the manuscript*.

## 9 Data availability

TopFracCCLE can be obtained as R-script at https://github.com/HuettelmaierLab/topFracCCLE. ENCODE accession IDs of the samples used in this study can be found in Table S4 and S5.

## References

[1] J. L. Almeida and C. T. Korch. Authentication of human and mouse cell lines by short tandem repeat (STR) DNA genotype analysis. In Assay Guidance Manual. Eli Lilly & Company and the National Center for Advancing Translational Sciences, Bethesda (MD), 2023.

[2] P. Hughes, D. Marshall, Y. Reid, H. Parkes, and C. Gelber. The costs of using unauthenticated, over-passaged cell lines: how much more data do we need? Biotechniques, 43(5):575, 577–8, 581–2 passim, November 2007.

[3] R. Barallon, S. R. Bauer, J. Butler, A. Capes-Davis, W. G. Dirks, E. Elmore, M. Furtado, M. C. Kline, A. Kohara, G. V. Los, R. A. MacLeod, J. R. Masters, M. Nardone, R. M. Nardone, R. W. Nims, P. J. Price, Y. A. Reid, J. Shewale, G. Sykes, A. F. Steuer, D. R. Storts, J. Thomson, Z. Taraporewala, C. Alston-Roberts, and L. Kerrigan. Recommendation of short tandem repeat profiling for authenticating human cell lines, stem cells, and tissues. In Vitro Cell Dev Biol Anim, 46(9):727–732, Oct 2010.

[4] E. Fasterius, C. Raso, S. Kennedy, N. Rauch, P. Lundin, W. Kolch, M. Uhlén, and C. Al-Khalili Szigyarto. A novel RNA sequencing data analysis method for cell line authentication. PLoS One, 12(2):e0171435, February 2017.

[5] Tabrez A Mohammad, Yun S Tsai, Safwa Ameer, Hung-I Harry Chen, Yu-Chiao Chiu, and Yidong Chen. CeL-ID: cell line identification using RNA-seq data. BMC Genomics, 20(Suppl 1):81, February 2019.

[6] J. Barretina, G. Caponigro, N. Stransky, K. Venkatesan, A. A. Margolin, S. Kim, C. J. Wilson, J. r, G. V. Kryukov, D. Sonkin, A. Reddy, M. Liu, L. Murray, M. F. Berger, J. E. Monahan, P. Morais, J. Meltzer, A. Korejwa, J. Valbuena, F. A. Mapa, J. Thibault, E. Bric-Furlong, P. Raman, A. Shipway, I. H. Engels, J. Cheng, G. K. Yu, J. Yu, P. Aspesi, M. de Silva, K. Jagtap, M. D. Jones, L. Wang, C. Hatton, E. Palescandolo, S. Gupta, S. Mahan, C. Sougnez, R. C. Onofrio, T. Liefeld, L. MacConaill, W. Winckler, M. Reich, N. Li, J. P. Mesirov, S. B. Gabriel, G. Getz, K. Ardlie, V. Chan, V. E. Myer, B. L. Weber, J. Porter, M. Warmuth, P. Finan, J. L. Harris, M. Meyerson, T. R. Golub, M. P. Morrissey, W. R. Sellers, R. Schlegel, and L. A. Garraway. The Cancer Cell Line Encyclopedia enables predictive modelling of anticancer drug sensitivity. Nature, 483(7391):603–607, Mar 2012.

[7] W. G. Dirks, R. A. F. MacLeod, Y. Nakamura, A. Kohara, Y. Reid, H. Milch, H. G. Drexler, and H. Mizusawa. Cell line cross-contamination initiative: an interactive reference database of STR profiles covering common cancer cell lines. Int. J. Cancer, 126(1):303–304, January 2010.

[8] A. Capes-Davis, Y. A. Reid, M. C. Kline, D. R. Storts, E. Strauss, W. G. Dirks, H. G. Drexler, R. A. F. MacLeod, G. Sykes, A. Kohara, Y. Nakamura, E. Elmore, R. W. Nims, C. Alston-Roberts, R. Barallon, G. V. Los, R. M. Nardone, P. J. Price, A. Steuer, J. Thomson, J. R. W. Masters, and L. Kerrigan. Match criteria for human cell line authentication: where do we draw the line? Int. J. Cancer, 132(11):2510–2519, June 2013.

[9] D. Kim, B. Langmead, and S. L. Salzberg. HISAT: a fast spliced aligner with low memory requirements. Nat Methods, 12(4):357–360, Apr 2015.

[10] H. Li, B. Handsaker, A. Wysoker, T. Fennell, J. Ruan, N. Homer, G. Marth, G. Abecasis, R. Durbin, and 1000 Genome Project Data Processing Sub-group. The sequence Alignment/Map format and SAMtools. Bioinformatics, 25(16):2078–2079, August 2009.

[11] Y. Liao, G. K. Smyth, and W. Shi. featureCounts: an efficient general purpose program for assigning sequence reads to genomic features. Bioinformatics, 30(7):923–930, Apr 2014.

[12] B. L. Aken, P. Achuthan, W. Akanni, M. R. Amode, F. Bernsdorff, J. Bhai, K. Billis, D. Carvalho-Silva, C. Cummins, P. Clapham, L. Gil, C. G. Girón, L. Gordon, T. Hourlier, S. E. Hunt, S. H. Janacek, T. Juettemann, S. Keenan, M. R. Laird, I. Lavidas, T. Maurel, W. McLaren, B. Moore, D. N. Murphy, R. Nag, V. Newman, M. Nuhn, C. K. Ong, A. Parker, M. Patricio, H. S. Riat, D. Sheppard, H. Sparrow, K. Taylor, A. Thormann, A. Vullo, B. Walts, S. P. Wilder, A. Zadissa, M. Kostadima, F. J. Martin, M. Muffato, E. Perry, M. Ruffier, D. M. Staines, S. J. Trevanion, F. Cunningham, A. Yates, D. R. Zerbino, and P. Flicek. Ensembl 2017. Nucleic Acids Res, 45(D1):D635–D642, 01 2017.

[13] A. R. Quinlan. BEDTools: The swiss-army tool for genome feature analysis. Curr. Protoc. Bioinformatics, 47(1):11.12.1–34, September 2014.

[14] P. Danecek, J. K. Bonfield, J. Liddle, J. Marshall, V. Ohan, M. O. Pollard, A. Whitwham, T. Keane, S. A. McCarthy, R. M. Davies, and H. Li. Twelve years of SAMtools and BCFtools. GigaScience, 10(2), 02 2021. giab008.

[15] R Core Team. R: A Language and Environment for Statistical Computing. R Foundation for Statistical Computing, Vienna, Austria, 2024.

[16] M. Kuhn. Building predictive models in r using the caret package. Journal of Statistical Software, 28(5):1–26, 2008.

[17] E. Pasolli, L. Schiffer, P. Manghi, A. Renson, V. Obenchain, D. T. Truong, F. Beghini, F. Malik, M. Ramos, J. B. Dowd, C. Huttenhower, M. Morgan, N. Segata, and L. Waldron. Accessible, curated metagenomic data through ExperimentHub. Nat. Methods, 14(11):1023–1024, October 2017.

[18] Hadley W. ggplot2: Elegant Graphics for Data Analysis. Springer-Verlag New York, 2016.

[19] A. Kozomara, M. Birgaoanu, and S. Griffiths-Jones. miRBase: from microRNA sequences to function. Nucleic Acids Res., 47(D1):D155–D162, January 2019.

[20] Y. Luo, B. C. Hitz, I. Gabdank, J. A. Hilton, M. S. Kagda, B. Lam, Z. Myers, P. Sud, J. Jou, K. Lin, U. K. Baymuradov, K. Graham, C. Litton, S. R. Miyasato, J. S. Strattan, O. Jolanki, J-W. Lee, F. Y. Tanaka, P. Adenekan, E. O’Neill, and J. M. Cherry. New developments on the encyclopedia of DNA elements (ENCODE) data portal. Nucleic Acids Res., 48(D1):D882–D889, January 2020.

[21] J. C. Melms, S. Vallabhaneni, C. E. Mills, C. Yapp, J-Y. Chen, E. Morelli, P. Waszyk, S. Kumar, D. Deming, N. Moret, S. Rodriguez, K. Subramanian, M. Rogava, A. N. R. Cartwright, A. Luoma, S. Mei, T. J. Brinker, D. M. Miller, A. Spektor, D. Schadendorf, N. Riggi, K. W. Wucherpfennig, P. K. Sorger, and B. Izar. Inhibition of haspin kinase promotes cell-intrinsic and extrinsic antitumor activity. Cancer Res., 80(4):798–810, February 2020.

[22] A. Bairoch. The cellosaurus, a cell-line knowledge resource. J. Biomol. Tech., 29(2):25–38, July 2018.

