## supplemental figures for "RNA-seq derived sequence variations are excellent features for cell line identification"

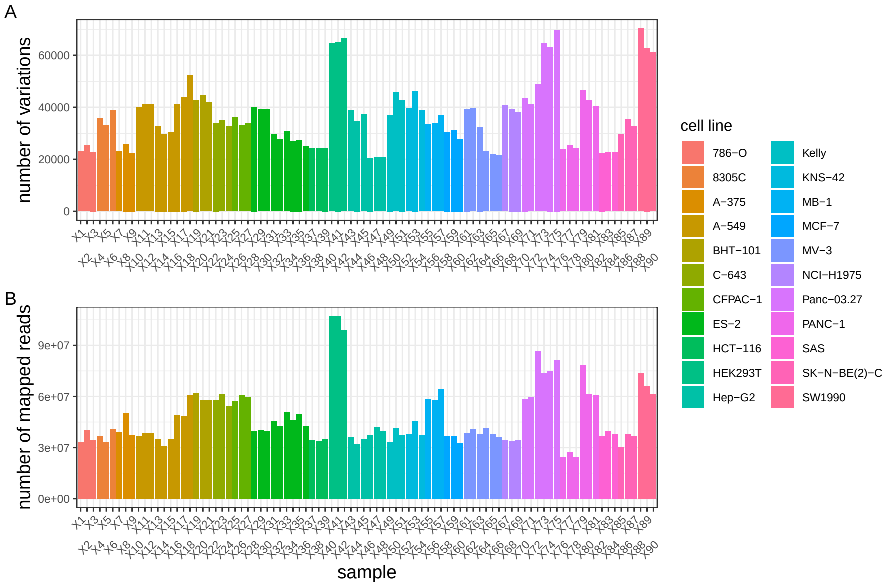


Figure S1: Variant calling and sequencing depth variations. **A** Number of detected sequence variations per sample compared to the reference genome (UCSC hg38). **B** Number of sequencing reads mapped to the reference genome (UCSC hg38) per sample.

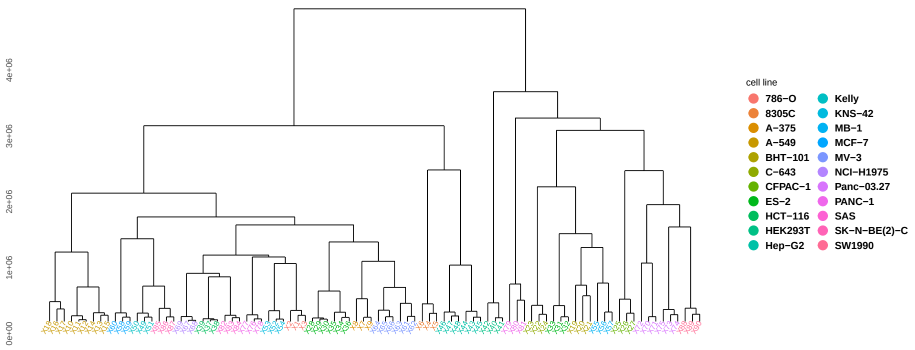


Figure S2: Hierarchical clustering of cell lines samples based on FPKM values

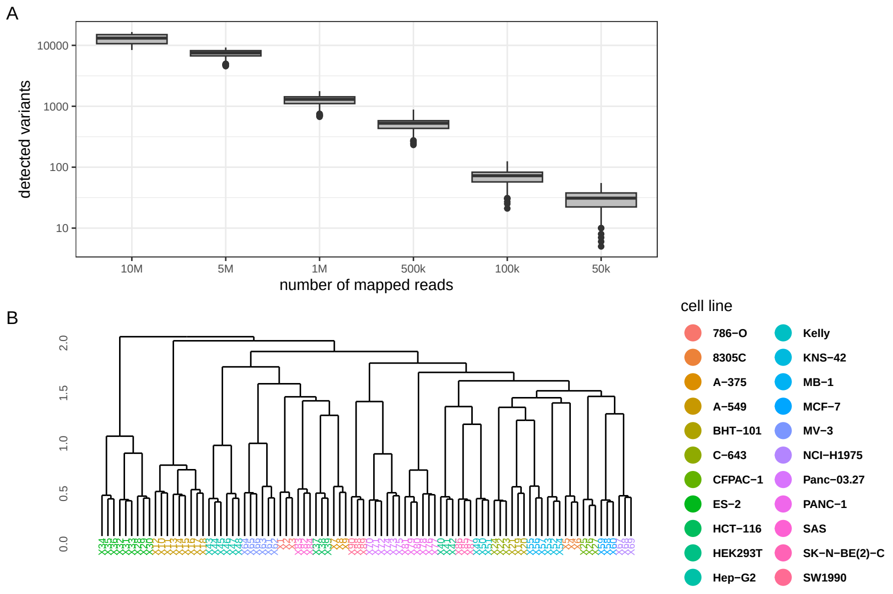


Figure S3: Clustering of cell lines by poly-A enriched RNAseq-derived sequence variations. **A** Detected sequence variations in dependence of sequencing reads. **B** Hierarchical clustering of samples based on sequence variations detected by randomly selecting 500,000 reads per sample.

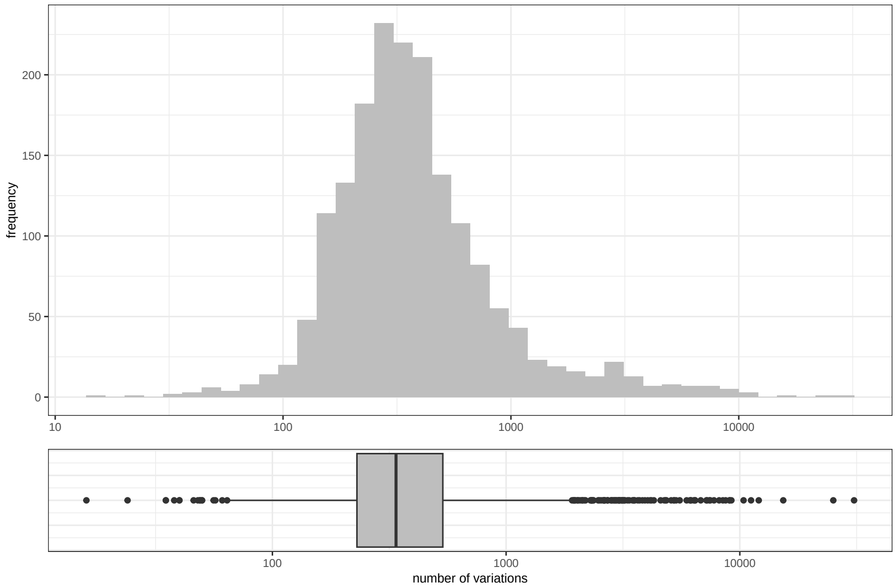


Figure S4: Distribution of reported SNP numbers in distinct cell lines obtained from the CCLE.

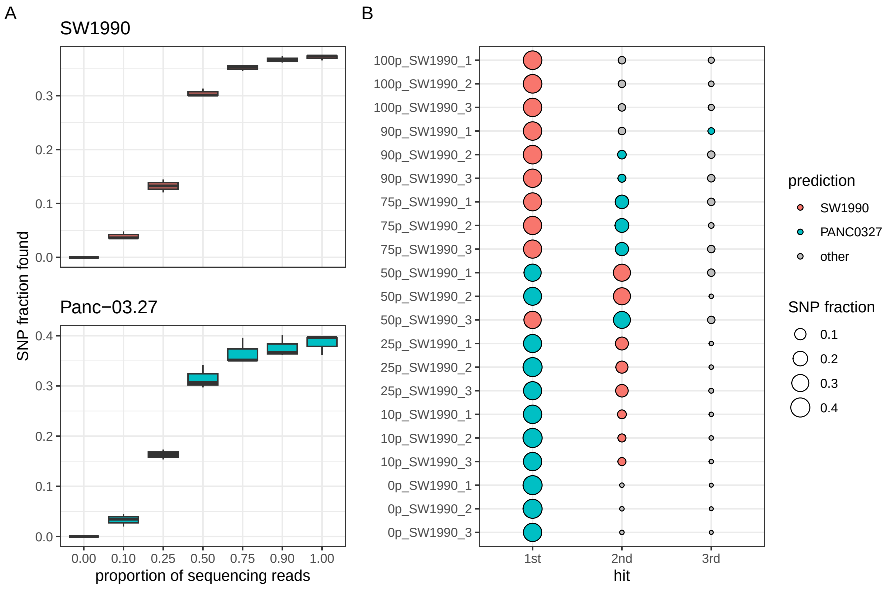


Figure S5: TopFracCCLE predictions on in silico contamination simulations **A** Proportion of detected SNPS in dependence of the relative abundance of the specified cell line among sequencing reads. **B** Top three prediction results of the indicated samples. Numbers of the sample labels specify the proportion of the respective cell line. Proportion of the other cell line equals 100 minus the indicated number.
